# A distinct brain expression profile for genes within large introgression deserts and under positive selection in *Homo sapiens*

**DOI:** 10.1101/2021.03.26.437167

**Authors:** Raül Buisan, Juan Moriano, Alejandro Andirkó, Cedric Boeckx

## Abstract

Analyses of ancient DNA from extinct hominins have provided unique insights into the complex evolutionary history of *Homo sapiens*, intricately related to that of the Neanderthals and the Denisovans as revealed by several instances of admixture events. These analyses have also allowed the identification of introgression deserts: genomic regions in our species that are depleted of ‘archaic’ haplotypes. The presence of genes like *FOXP2* in these deserts has been taken to be suggestive of brain-related functional differences between *Homo* species. Here, we seek a deeper characterization of these regions, taking into account signals of positive selection in our lineage. Analyzing publicly available transcriptomic data from the human brain at different developmental stages, we found that structures outside the cerebral neocortex, in particular the cerebellum, the striatum and the mediodorsal nucleus of the thalamus show the most divergent transcriptomic profiles when considering genes within large introgression deserts and under positive selection.

## 1 Introduction

The availability of high-coverage genomes from our closest extinct relatives, the Neanderthals and Denisovans, constitutes a significant advance in the range of questions one can ask about the deep history of our species [1–4]. One of the main themes emerging from this progress is interbreeding. In recent years, a fairly large number of admixture events between Neanderthals, Denisovans and Sapiens populations have been postulated. A recent review [5] considers that at least four such events are supported by strong evidence.

While it is important to ask whether our species benefited from these admixture events (so-called adaptive introgression, where alleles inherited from other hominins rose to high frequency as a result of positive selection after gene flow), it is also worth examining regions of the genomes that are depleted of alleles resulting from gene flow from other hominins [6–10]. Such regions are called introgression deserts (sometimes also ‘genomic islands of divergence/speciation’ [11]).

There are multiple reasons why genetic differences that arose after the divergence of populations may not be well tolerated [12]: there could be negative selection on ‘archaic’ variants (deleterious changes on the ‘archaic’ lineage), or positive selection on human-specific variants (adaptive changes on the human lineage), or it may be due to drift. It is reasonable to expect, and indeed has been shown, that the X chromosome constitutes such a desertic region (not only in our species [13, 14]). This could be due to repeated selective sweeps on this chromosome: genes involved in reproduction on this chromosome might act as strong reproductive barriers between populations [15].

In the case of modern humans, other genomic regions are devoid of Neanderthal and Denisovan introgression, for reasons that are perhaps less obvious, and therefore worth investigating further. A recent study [8] identifies four large deserts depleted of Neanderthal introgression, partially overlapping with a previous independent study [7]. As pointed out in [12, 16], since there quite possibly were several different pulses of gene flow between us and our closest relatives [17], the depletion observed in these four regions must have been reinforced repeatedly, and given the size of the deserts, it is reasonable that the ‘archaic’ haplotype was purged within a short time after the gene flow event (as predicted by mathematical modeling on whole-genome simulations [18] and as evidenced in the analysis of genome-wide data from the earliest Late Pleistocene modern humans known to have been recovered in Europe [19]).

The presence of *FOXP2*, a gene known for its role in language [20, 21], in one of these large deserts has attracted attention [16], as it raises the possibility that the incompatibility between Sapiens and other hominin in such persistent introgression deserts may point to (subtle, but real) cognitive differences. Indeed, the presence in such deserts of not only *FOXP2* but also other genes like *ROBO1, ROBO2*, and *EPHA3*, all independently associated with language traits [22–24], together with an earlier observation that genes within large deserts are significantly enriched in the developing cerebral cortex and in the adult striatum [7], suggest a possible point of entry into some of the most distinctive aspects of the human condition [25]. Such considerations, combined with independent evidence that introgressed Neanderthal alleles show significant downregulation in brain regions [26], motivated us to focus on the brain in this study.

Specifically, we investigated the four largest genomic regions that resisted ‘archaic’ introgression reported in [8] jointly with the most comprehensive signals of positive selection in our lineage [27], a combination that, to our knowledge, has not been previously studied in detail. We sought to characterize the expression profile of genes within our regions of interest by analyzing transcriptomic data from several brain regions at different developmental stages from [28], which allows for greater resolution than the Allen Brain Atlas data used in [7], especially for early stages of development. Three of the brain regions under study showed marked transcriptomic divergence (as assessed via pairwise comparisons of their Euclidean distances based on statistically significant principal components): the cerebellum, the striatum and the thalamus. Among the genes at the intersection of regions under positive selection and large deserts of introgression, we found *CADPS2, ROBO2*, or *SYT6*, involved in neurotrophin release, axon guidance and neuronal proliferation.

## 2 Results

### 2.1 Genes in large deserts of introgression have different expression levels relative to the rest of the genome

We set to understand whether the mean expression of genes in large deserts of introgression [8] and the positively selected regions within them (extracted from [27]) is significantly different compared to the rest of the genome, using publicly available transcriptomic data from the human brain [28]. To this end, we selected random regions of the genome (*n* = 1000), excluding the large deserts, of the same average length (i.e., 15 million base-pairs), with a possible deviation of 1 million base-pairs to account for the length variability between different deserts of introgression. To avoid genomic regions with low genetic density that might skew the results, the randomized areas were required to hold at least as many genes (265) as the desertic regions reported in [8].

The mean expression of genes lying in random regions of the genome was summarized for each brain structure (and log2-transformed). A repeated-measures two-way anova shows that the mean expression of both sets of these regions is significantly different from the rest of the genome (*p* < 0.01 for both sets). A post-hoc pairwise anova (with Bonferroni correction) shows the difference between a gene expression value in a brain region as derived from the control set and that obtained from the genes in our two sets of interest is significant for most structures. An outlier’s Grubbs test shows that the structures with the highest and lowest mean gene expression values in large deserts of introgression and the positively-selected windows within them fall inside the expected range of variability given the data (*p* > 0.01).

### 2.2 The cerebellar cortex, the striatum and the thalamus show divergent transcriptomic profiles when considering genes within large deserts of introgression and under positive selection

We then investigated the temporal progression of the expression of genes within large deserts of introgression and putative positively-selected regions analyzing RNA-seq data of different human brain regions at different developmental stages [28]. We found that the median expression of genes within large deserts and positively-selected regions is higher than those present in deserts alone, the former peaking at prenatal stages in neocortical areas and decreasing later on. Outside the cerebral neocortex, this pronounced prenatal peak is not observed and, specifically for the cerebellar cortex, the expression profile of these genes increases before birth and reaches the highest median expression from childhood to adulthood in comparison to the rest of structures (see Figure1A and Figure S1).

**Figure 1:**
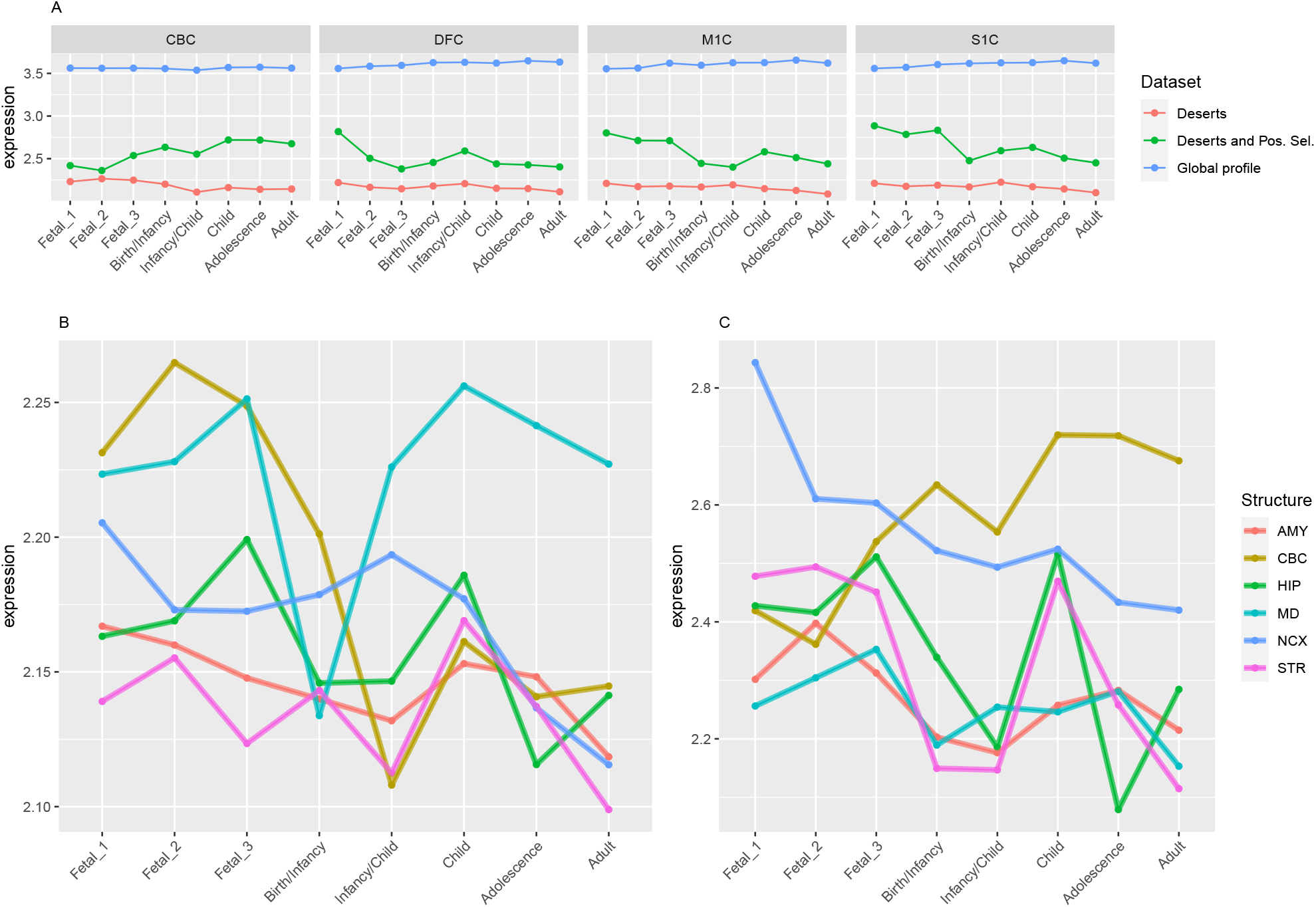
Median expression of genes in large deserts and deserts/positively-selected regions. A) Selected examples comparing genes in the global dataset (n=9358 genes; requiring median > 2), in large deserts of introgression (n=255) and in both deserts and positively-selected regions (n=12). A panel with all structures analyzed in this study is shown in Figure S1. In neocortical areas, genes within deserts and positively-selected regions peak at prenatal stages and decrease later, whereas the opposite pattern is found for the cerebellar cortex (see also C). Of note, the global dataset was filtered by setting a threshold of median expression higher than 2 (following [28]), to avoid genes with extremely low expression, while we kept any outlier for the two other datasets. B) Median expression profile of genes within large deserts. The cerebellar cortex, prenatally, and the mediodorsal nucleus of the thalamus prenatally and postnatally present the highest expression. C) Median expression profile of genes within deserts and positively-selected regions. Genes expressed in neocortical areas reach the highest expression at the early fetal stages, whereas the cerebellarcortex, from birth until adulthood, remains with the highest expression profile. *Structures:* AMY, amygdala; CBC, cerebellar cortex; HIP, hippocampus; MD, mediodorsal nucleus of thalamus; NCX, neocortex; STR, striatum. *Stages:* Fetal 1: 12-13 post conception weeks (PCWs); Fetal 2: 16-18 PCW; Fetal 3: 19-22 PCW; Birth - Infancy: 35-37 PCW and 0-0.3 years; Infancy - Child: 0.5-2.5 years; Childhood: 2.8-10.7 years, Adolescence: 13-19 years; Adulthood: 21-64 years (as in [28])

In order to statistically evaluate the differences observed for each structure and developmental stage (see Figure1B,C), we performed a Principal Component Analysis and calculated the pairwise Euclidean distances between brain regions for each developmental stage using statistically significant principal components (*p* < 0.05) as assessed via the JackStraw analysis implemented in Seurat [29]. For genes within large deserts of introgression overlapping putative positively-selected regions, we performed dimensionality reduction on the first two principal components. Due to the low number of genes at this intersection (n=12), the second principal component did not report statistical significance. The sum of the percentage of variance explained by first and second components is around 50%.

For genes that reside in the deserts of introgression under consideration, the cerebellum stand out as the structure with the most divergent transcriptomic profile at postnatal stages, from childhood to adulthood (after stage-specific pairwise comparisons; Wilcoxon rank sum test with Bonferroni correction; significance was considered if *p* < .01; see Figure2). For genes under positive selection that are also found within introgression deserts, the cerebellum still remains as the most transcriptomically divergent structure postnatally (birth/infancy, childhood, adolescence and adulthood; see the caption of Figure 1 for the specific time points associated to each developmental stage). Moreover, prenatally, the cerebellum again (fetal stages 1 and 2) and the mediodorsal nucleus of the thalamus (fetal stage 1; see Suppl. Fig. S4,9) report the most significant differences in the pairwise comparisons. Previous research found that genes within large deserts are over-represented in the striatum at adolescence and adult stages [7]. In consonance with this finding, we found that the transcriptomic profile of the striatum for genes within large deserts is significantly different at adolescence and adulthood but also at fetal stage 3, while for genes within deserts under putative positive selection, significant differences are found at infancy and adolescence (see Suppl. Fig. S3,4,8).

**Figure 2:**
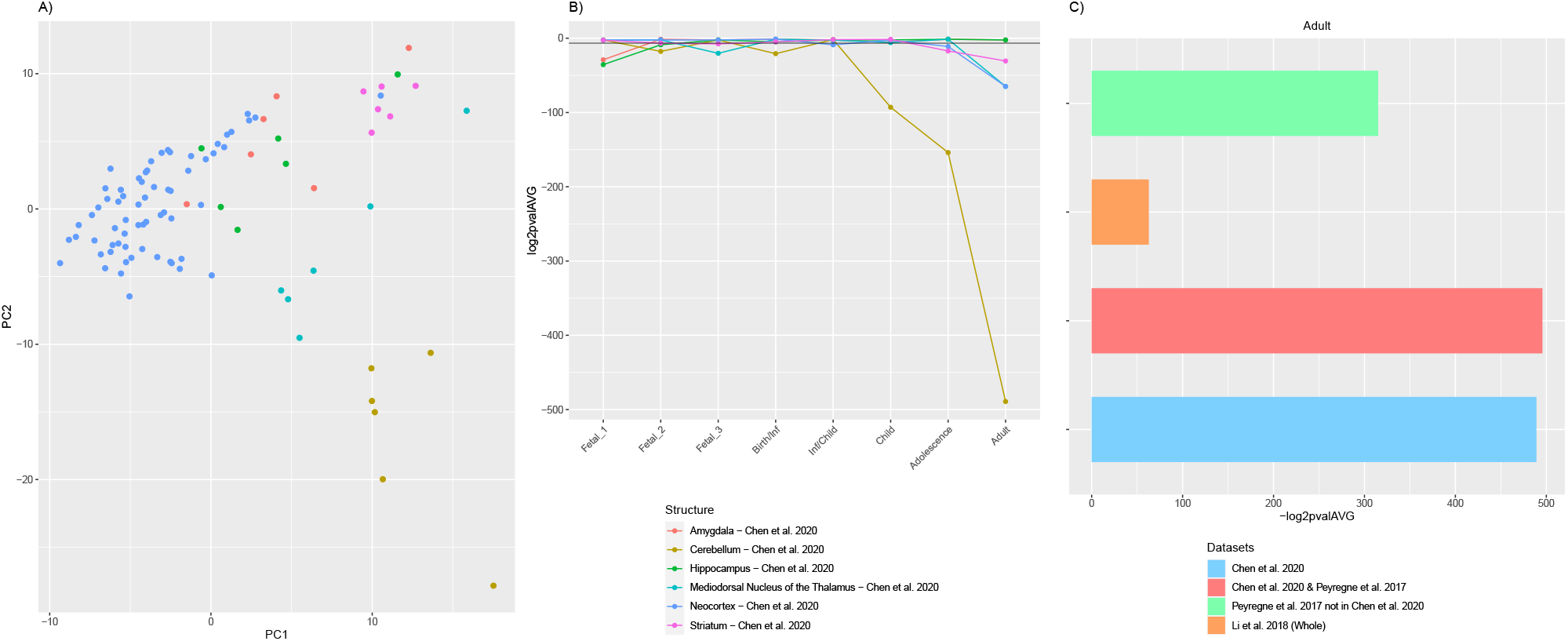
The cerebellum’s transcriptomic profile significantly diverges at postnatal stages. For genes within large deserts of introgression, the cerebellum reports the most significant differences when evaluating pairwise distances between structures based on the statistically significant principal components. A) Distribution of structures at the adult stage using the first two principal components. B) P-values (log2-transformed) obtained from pairwise comparisons among structures at each developmental stage (Wilcoxon rank sum test with Bonferroni correction). C) Contribution of genes within large deserts of introgression [8], deserts of introgression under putative positive selection [8, 27], regions under putative selection [27] not within large deserts, and the raw dataset used in this study [28] to the observed divergence at adult stage for the cerebellum, with the greatest value for genes within large deserts.

A similar profile postnatally obtains for the cerebellum when subsetting for genes under positive selection not present within large introgression deserts (marked differences form adulthood to childhood; see Suppl. Fig. S6). When evaluating the global expression profile (n=9358 genes), the cerebellum shows statistically significant differences also at postanatal stages (birth, infancy, childhood and adulthood) and the mediodorsal nucleus of the thalamus at fetal stage 3 and adulthood (see Suppl Fig. S5). All p-values can be found in Supplementary Material.

The trajectories of expression across developmental stages in genes within large deserts of introgression might be affected by positive selection. To control for this, we analyzed the contrast between a control group of genes not under positive selection but within deserts of introgression compared to those under positive selection in these same regions. We found that, within large deserts of introgression, genes under positive selection have an overall lower expression than those in regions not under positive selection (*p* = 0.0007, Kruskal-Wallis test). A linear regression model predicts that this effect is not structure-specific (p = 0.655), and that overall variability in the data is not explained by between-structure differences (*p* = 0.9904, anova test between fitted models that do and do not include brain regions as a variable). Expression linked to specific developmental stages diverges significantly between genes under positive selection and those that are not (0.0001, linear regression). However, a post-hoc Tukey test (corrected for repeated measures, Fig. S2) reveals that this difference holds only at the fetal stages. In portions of large deserts not under selection, the fetal period of development is significantly different from most posterior stages, while in genes under selective pressures only the first fetal stage is significantly different from post-fetal stages (with a significance threshold of *p* < 0.05).

### 2.3 Gene-specific expression trajectories of genes in the overlapping desertic and positively-selected regions

Focusing on the specific trajectories of genes at the intersection of large deserts and positively-selected regions (n=12 genes; Figure S1), we performed a segmented regression analysis (using the Trendy package [30]) filtering out genes with an adjusted *R*^2^ less than 0.5. As our analysis showed a marked increase of transcriptomic divergence at different developmental stages for the cerebellum, the striatum and the mediodorsal nucleus of the thalamus, we decided to focus on these structures.

For the cerebellum, *CADPS2* expression is the one that most closely mimics the observed pattern, with highest postnatal expression and a marked increased of its expression around birth and infancy (*R*^2^ 0.56; see Figure S7 and S10). This Ca^2+^-dependent activator protein is known to regulate exocytosis in granule cells, particularly neurotrophic factors BDNF and NT-3 release, and its knockout disrupts normal cerebellar development and causes an autistic-like behavioral phenotype in mice [31, 32]. In addition, decreasing expression through developmental stages was also found for *SYT6* and *ROBO2* (*R*^2^ 0.76 and 0.60, respectively; see Figure S10). Two other genes, *KCND2* and *ST7*, with comparatively high expression postnatally, did not pass however the adjusted *R*^2^ threshold (Figure S7). Regarding the thalamus, two genes within the overlapping desertic and positively-selected regions could be fitted with an adjusted *R*^2^ higher than 0.5: *ROBO2* and *ST7*. Both genes show higher expressions at prenatal stages, followed by a steady decline at around birth (*R*^2^ 0.65 and 0.61, respectively; see Figure S11). The roles of Robo2 in the thalamus have been studied as a receptor of the Slit/Robo signaling pathway which is critically involved in axon guidance. Indeed, Robo2 is highly expressed in the dorsal thalamus and cerebral cortex in the embryonic mouse brain and, in cooperation with Robo1, is required for the proper development of cortical and thalamic axonal projections [33].

Lastly, for the striatum, three genes within the overlapping desertic and positively-selected regions could be fitted with an adjusted *R*^2^ higher than 0.5. ST7, *ROBO2* and *SYT6* follow a V-shape profile with higher expression at prenatal stages, a decrease around birth, and increasing levels during later postnatal stages (*R*^2^ = 0.75, 0.57, and 0.53, respectively; see Figure S12). While the role of *ST7* in neurodevelopment remains to be elucidated, Robo2 is a receptor of the Slit/Robo signaling pathway which is critically involved in axon guidance [33], but also in the proliferation and differentiation of neural progenitors with possible different roles in dorsal and ventral telencephalon [34, 35]. *Syt6* is another synapse-related gene expressed in the developing basal ganglia [36], and in fact linked to the distinctive expression profile of this structure [37]. Additionally, *Syt6* shows a similar expression profile in the cerebellar cortex although at lower levels (see Figure S10), where is differentially expressed in *Cadps2* knockout mice [38].

## 3 Discussion

The current study brings out two main findings: the importance of structures beyond the cerebral neocortex in the attempt to characterize some of the most derived features of our species’ brain, and the fact that some of the strongest effect in these regions takes place at early stages of development. In this way our work adduces complementary evidence for the perinatal globularization phase as a species-specific ontogenic innovation [39],and also provides new evidence for the claim that brain regions outside the neocortex (cerebellum, thalamus, striatum) significantly contribute to this phenotype [26, 40–43].

To our knowledge this is the first study to characterize the effect of the cerebellum in the context of large introgression deserts. For the striatum, previous studies have already highlighted the relevance of this structure: genes carrying Neanderthal-derived changes and expressed in the striatum during adolescence exhibit a higher McDonald-Kreitman ratio [4]. In addition, using a different range of introgressed regions, it had already been noted [7] that genes within large deserts are significantly enriched in the striatum at adolescence and adult stages, which matches the life stages highlighted from our analysis using the most recent report of genomic regions depleted of archaic variants [8].

Naturally, the functional effects of these divergent developmental profiles for the cerebellum, the prenatal thalamus or the striatum remain to be understood, particularly in the context of the possible *Homo*-divergent regulation of the genes highlighted in this study. This is especially relevant in light of emerging evidence that selection against DNA introgression is stronger in regulatory regions [44], which in addition have been found to be over-represented in putative positively-selected regions in *Homo sapiens* [27, 45]. But the fact that early developmental stages are key holds the promise of using brain organoid technology to probe the nature of these differences, since such *in vitro* techniques best track these earliest developmental windows [46–48].

The fact that FOXP2 expression is known to be particularly high in the brain regions highlighted here and in complementary studies [49] may help shed light on why *FOXP2* is found in introgression deserts in modern human genomes. As pointed out in [16], this portion of chromosome 7 is not a desert for introgression in other great apes, nor did it act as a barrier for gene flow from Sapiens into Neanderthals. As such, it may indeed capture something genuinely specific about our species.

## 4 Methods

Genes within large deserts of introgression or putative positively-selected regions were obtained via BioMart R package [50], using the respective genomic region coordinates as input and filtering by protein-coding genes.

### mRNA-seq analysis

Publicly available transcriptomic data of the human brain at different developmental stages was retrieved from [28]. Reads per kilo base per million mapped reads (RPKM) normalized counts were log-transformed and then subsetted to select genes either in large deserts of introgression or in both deserts and putative positively-selected regions. The complete log-transformed, RPKM normalized count matrix was subsetted to select genes with median expression value > 2, as in [28], while no median filtering was employed for the subsets of genes within deserts and positively-selected regions, due to the potential relevance of the outliers in these specific regions for the purposes of our study. To assess transcriptomic variability between brain regions accounted for by genes either in large deserts or in deserts and positively-selected regions, we performed principal component analysis and calculated the pairwise Euclidean distances between brain regions for each dataset (following [28]). We then statistically evaluated such differences at each developmental stage using pairwise Wilcoxon tests with Bonferroni correction. Significant differences were considered if *p* < .01.

### Gene-specific expression trajectories

The R package Trendy [30] was used to perform segmented regression analysis and characterize the expression trajectories of genes within both deserts of introgression and putative positively-selected regions (12 genes). The normalized RPKM values (from [28]) in the form of a gene-by-time samples matrix was used to fit each gene expression trajectory to an optimal segmented regression model. Genes were considered if their adjusted *R*^2^ was > 0.5. In addition, a maximum number of breakpoints (significant changes in gene expression trajectory) was set at 3, minimum number of samples in each segment at 2, and minimum mean expression, 2.

The permutation tests using gene expression data from [28] were done using the regioneR package [51] at *n* = 1000.

**Table 1:** Genomic coordinates used in this study.

| Set | Chromosome | Start | End |
| --- | --- | --- | --- |
| **Large deserts of introgression [8]** | 1 | 105400000 | 120600000 |
|  | 3 | 74100000 | 89300000 |
|  | 7 | 106200000 | 123200000 |
|  | 8 | 49400000 | 66500000 |
| **Regions under positive selection [27] within deserts** | 1 | 113427676 | 113560554 |
|  | 1 | 114641362 | 114645248 |
|  | 1 | 119322276 | 119387279 |
|  | 3 | 77027847 | 77034264 |
|  | 7 | 106877730 | 107233808 |
|  | 7 | 116762909 | 116773234 |
|  | 7 | 120147456 | 120174406 |
|  | 7 | 122320035 | 122406480 |

## Data and material availability

Datasets and code are made available at https://github.com/jjaa-mp/desertsHomo.

## Author Contributions

Conceptualization: CB & AA & JM & RGB; Data Curation: AA & JM & RGB; Formal Analysis: AA & JM & RGB; Visualization: CB & AA & JM & RGB; Writing – Original Draft Preparation: CB & AA & JM & RGB; Writing – Review & Editing: CB & AA & JM & RGB; Supervision: CB; Funding Acquisition: CB.

## Funding statement

CB acknowledges support from the Spanish Ministry of Economy and Competitiveness (grant PID2019-107042GB-I00), MEXT/JSPS Grant-in-Aid for Scientific Research on Innovative Areas #4903 (Evolinguistics: JP17H06379), Generalitat de Catalunya (2017-SGR-341), and the Foundation BBVA (Leonardo fellowship). JM acknowledges financial support from the Departament d’Empresa i Coneixement, Generalitat de Catalunya (FI-SDUR 2020). AA acknowledges financial support from the Spanish Ministry of Economy and Competitiveness and the European Social Fund (BES-2017-080366).

## Notes

https://github.com/jjaa-mp/desertsHomo

### Competing Interest Statement

The authors have declared no competing interest.

### Summary of Updates

Additional analyses in sections 2.2 and 2.3, additional Supplementary figures

