## Supplementary Figures for "A distinct brain expression profile for genes within large introgression deserts and under positive selection in *Homo sapiens*"

for

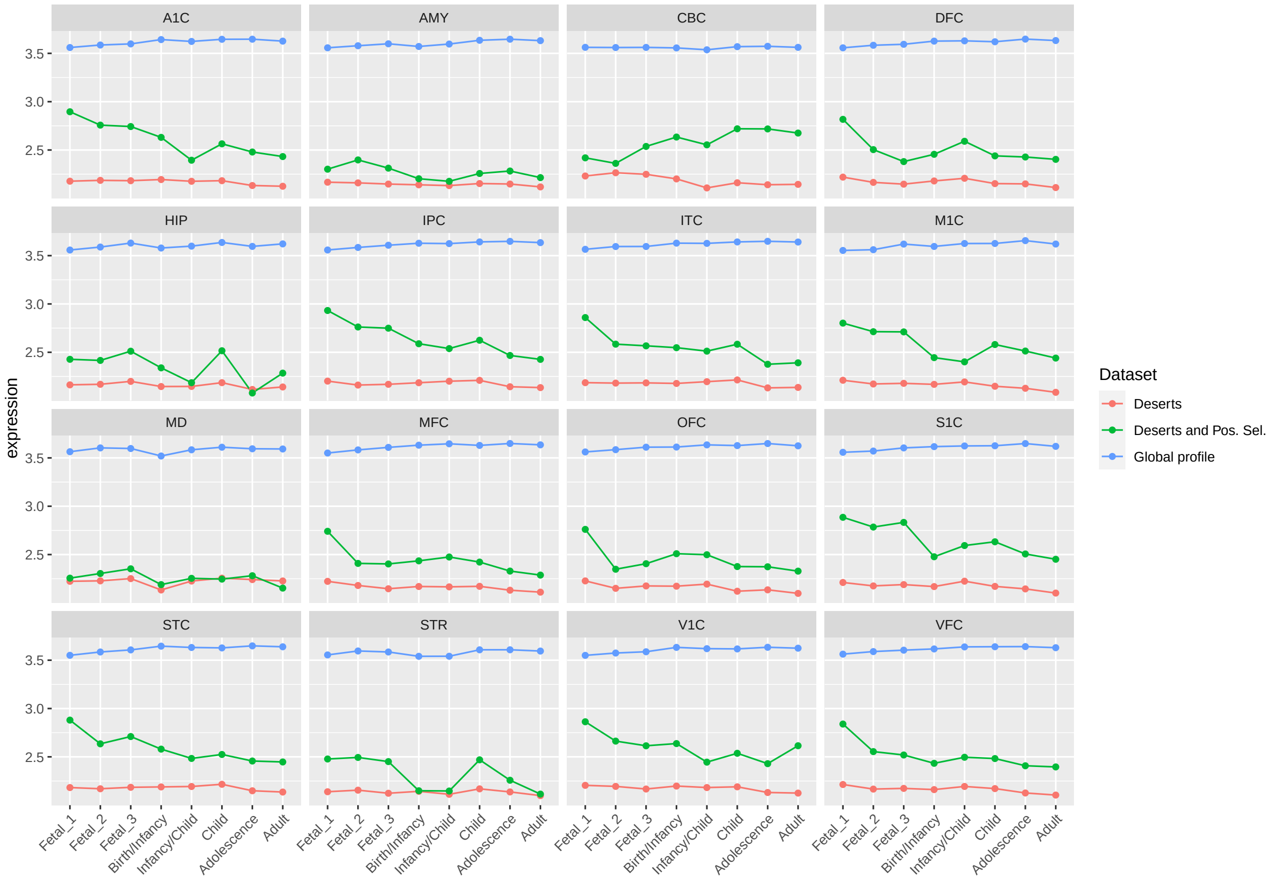


Figure 1: **Median expression profile of genes within large deserts, deserts/positively-selected regions and the global dataset, across structures and stages.** Related to Figure1, genes within deserts/positively-selected regions have higher expression in comparison to the broader subset of genes within deserts, with a peak of expression at prenatal stages in neocortical areas. This peak is not observed in non-neocortical areas and, in the case of the cerebellar cortex, the genes (within deserts/positively-selected regions) show increasing expression at postnatal stages.
A1C, primary auditory cortex; AMY, amygdala; CBC, cerebellar cortex; DFC, dorsolateral prefrontal cortex; HIP, hippocampus; IPC, posterior inferior parietal cortex; ITC, inferior temporal cortex; M1C, primary motor cortex; MD, mediodorsal nucleus of thalamus; MFC, medial prefrontal cortex; OFC, orbital prefrontal cortex; S1C, primary somatosensory cortex; STC, superior temporal cortex; STR, striatum; V1C, primary visual cortex; VFC, ventrolateral prefrontal cortex.

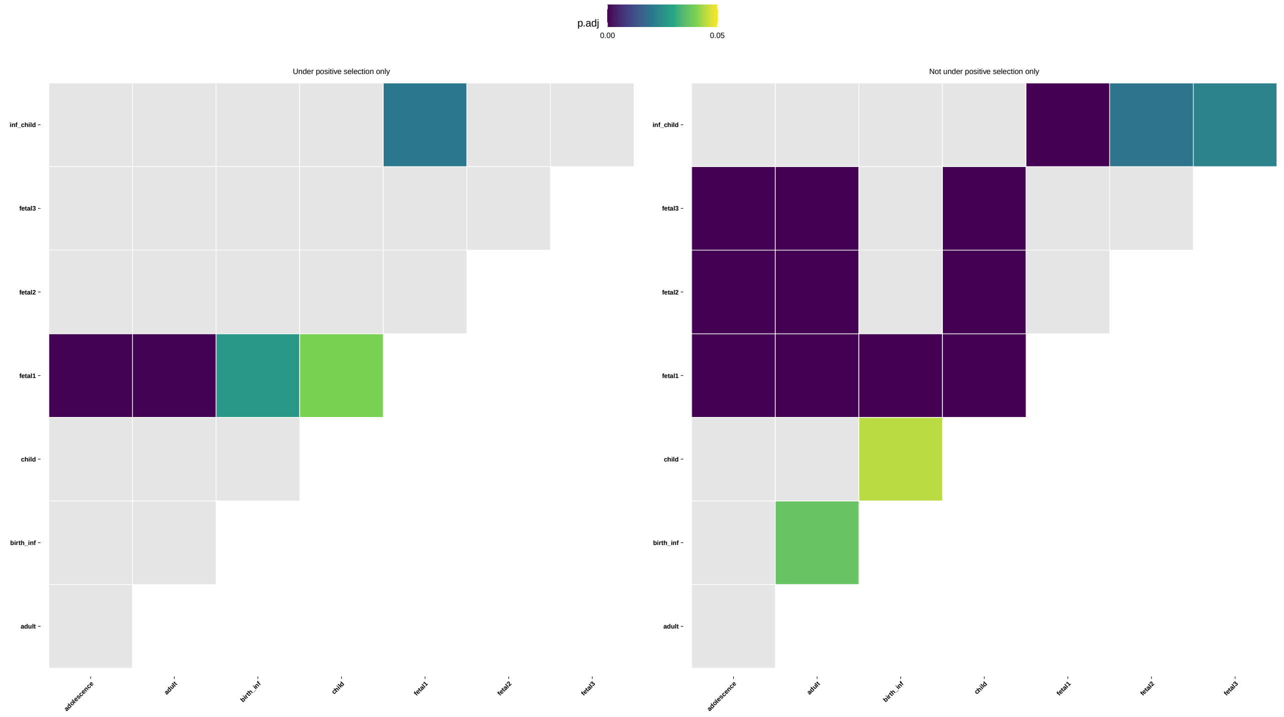


Figure 2: **Visualization of a pairwise post-hoc Tukey test** comparing the mean expression of genes in deserts of introgression under the effect of positive selection (left) and in genes not affected by positive selection (right) in each of the developmental stages included in [1].

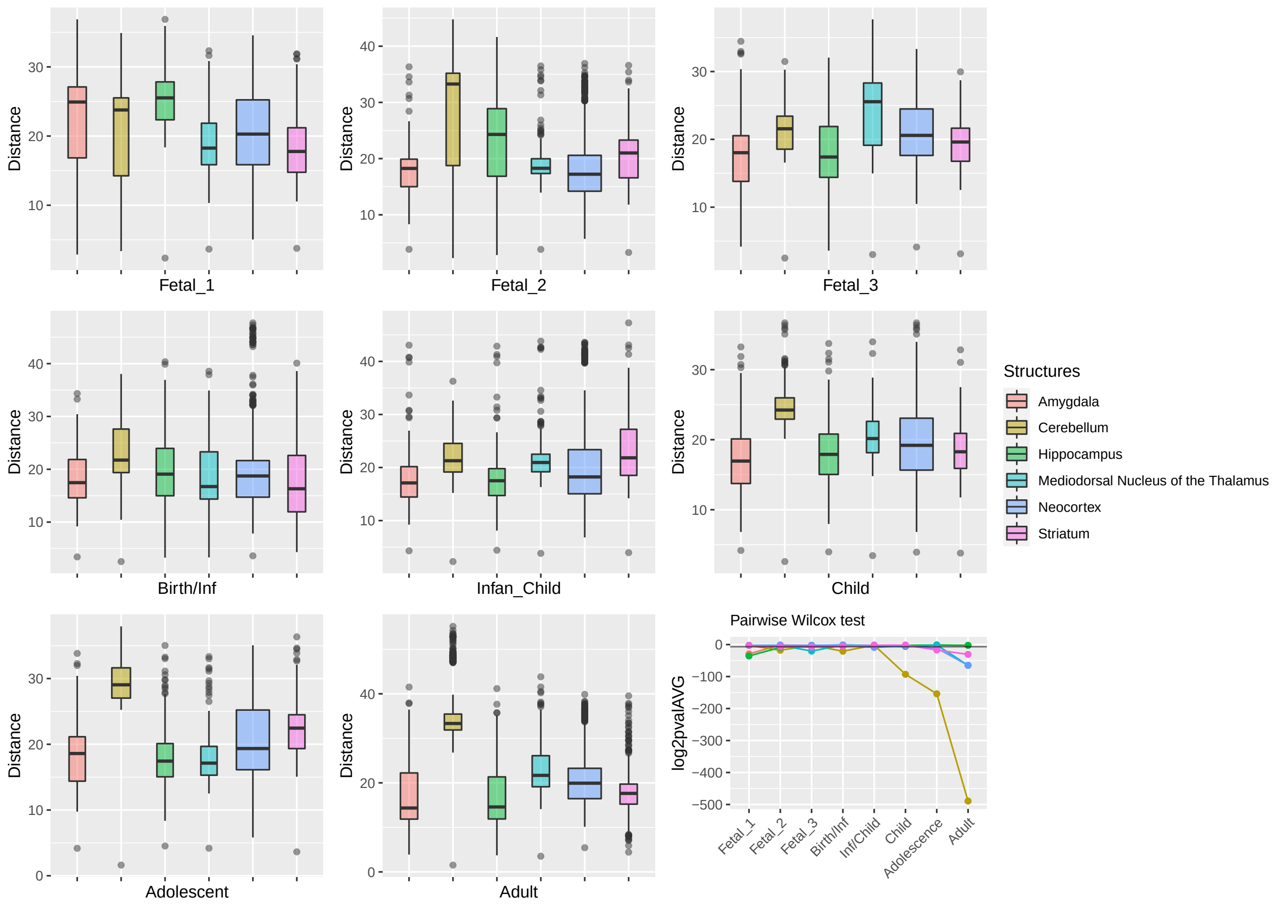


Figure 3: **Expression profile: Genes within large deserts.** One structure, the cerebellum, reports the most significant differences (pairwise Wilcoxon test with Bonferroni correction) encompassing postnatal stages: childhood ($p = 1.06\text{x}10^{-28}$), adolescence ($p = 4.71\text{x}10^{-47}$), adulthood ($p = 5.54\text{x}10^{-148}$); also at birth ($p = 5.64\text{x}10^{-7}$) and fetal stage 2 ($p = 4,79\text{x}10^{-6}$). Significant differences are also found for the thalamus (fetal stage 3 and adulthood), the hippocampus (fetal stages 1 and 2) or the striatum (fetal stage 3, adolescence and adulthood).
The boxplots show the values of the pairwise Euclidean distances at each stage for each structure. The line graph represents the average p-value (log2-transformed for representational purposes) for each pairwise comparison between structures at each stage. The horizontal black line denotes $p = 0.01$.

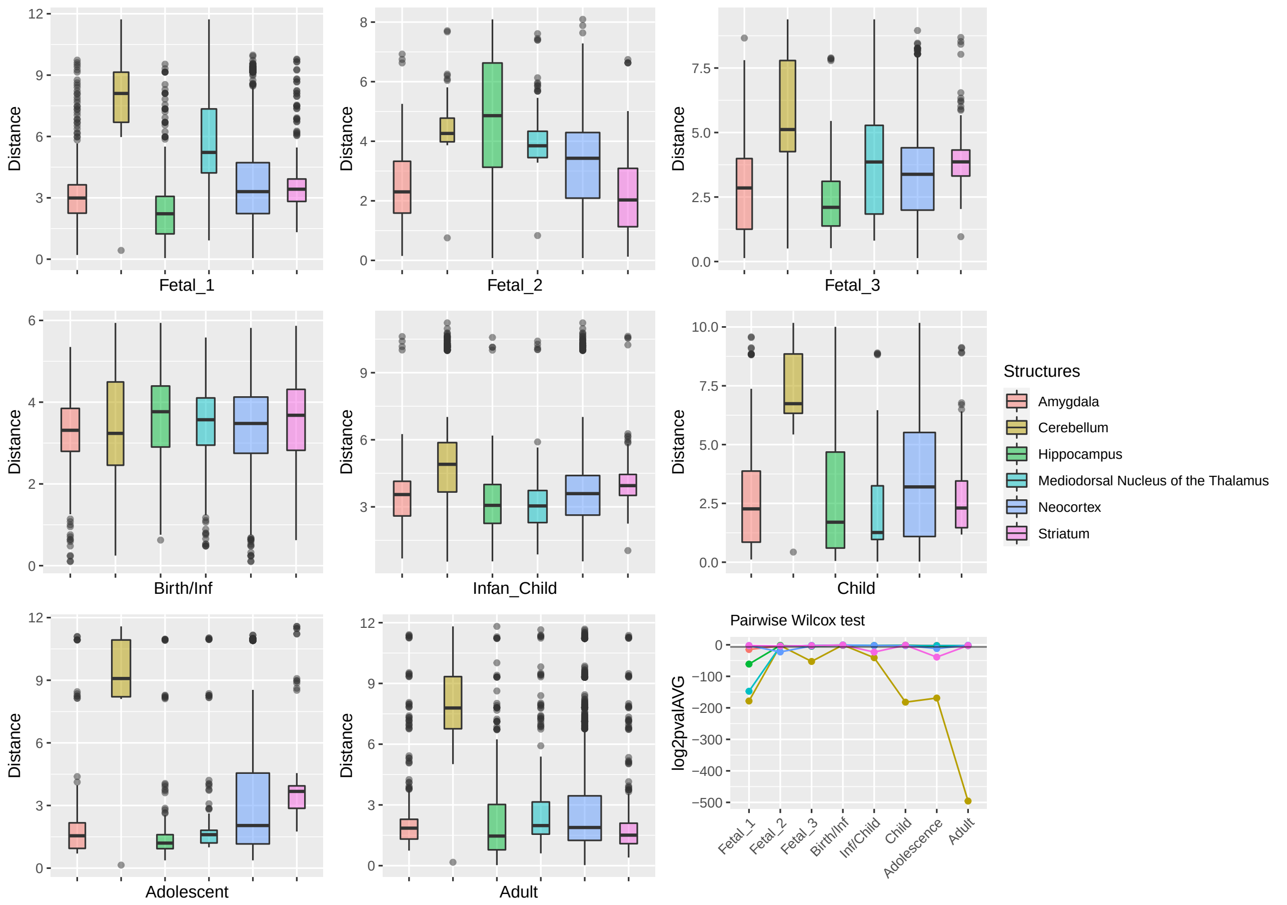


Figure 4: **Expression profile: Genes under positive selection within large deserts.** The two first principal components were selected to calculate the pairwise comparisons between structures at each stage (Wilcoxon test with Bonferroni correction). Significant results were obtained for the cerebellum more prominently at childhood $p = 1.64\text{x}10^{-55}$, adolescence $p = 1.45\text{x}10^{-51}$, and adulthood $p = 5.83\text{x}10^{-150}$. Prenatally the cerebellum and the thalamus show the most significant differences in comparison to the rest of structures, at fetal stage 1 ($p = 2.1\text{x}10^{54}$ and $p = 4.47\text{x}10^{-45}$, respectively). Other significant results are found at specific stages for the striatum (infancy and adolescence), neocortex (fetal stage 2), or the amygdala (fetal stages 1 and 3). The horizontal black line in the line plot denotes $p = 0.01$.

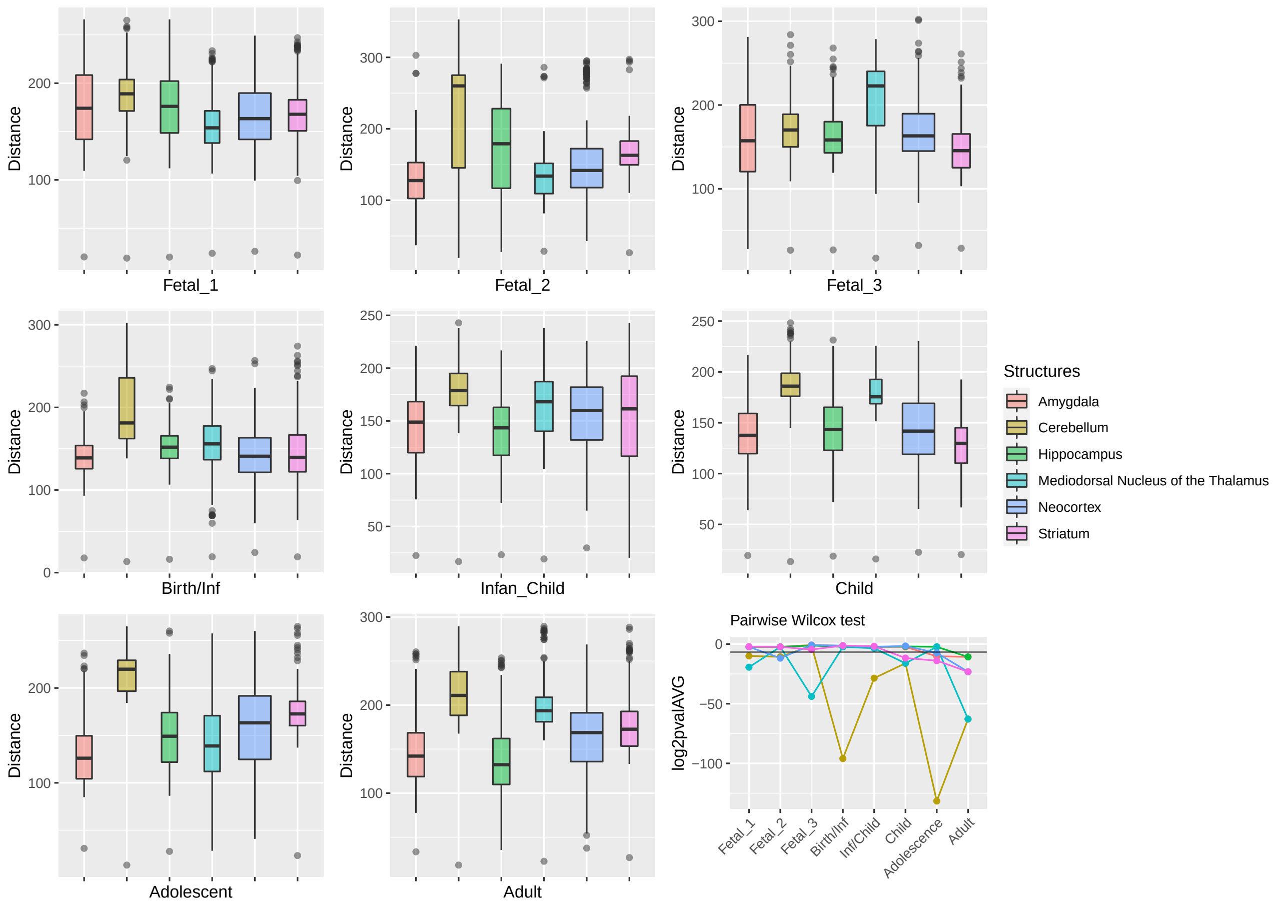


Figure 5: **Global profile of genes across stages and structures.** We processed the mRNA-seq data from [1] filtering out genes with a median across stages and structures less than 2, resulting in a total of 9358 genes. As described in section 4 of the main text, log-transformed, RPKM normalized counts were used to calculate pairwise Euclidean distances using statistically significant principal components for each stage. The most significant differences (pairwise Wilcoxon test with Bonferroni correction) are found for the cerebellum at postnatal stages (birth $p = 1.29\text{x}10^{-29}$; infancy $p = 2.53\text{x}10^{-9}$; adolescence $p = 2.44\text{x}10^{-30}$; adulthood $p = 1.21\text{x}10^{-19}$), and for the thalamus at fetal stage 3 ($p = 6.12\text{x}10^{-14}$) and adulthood ($p = 1.21\text{x}10^{-19}$) . All p-values can be found in Supplementary Material. The horizontal black line in the line plot denotes $p = 0.01$.

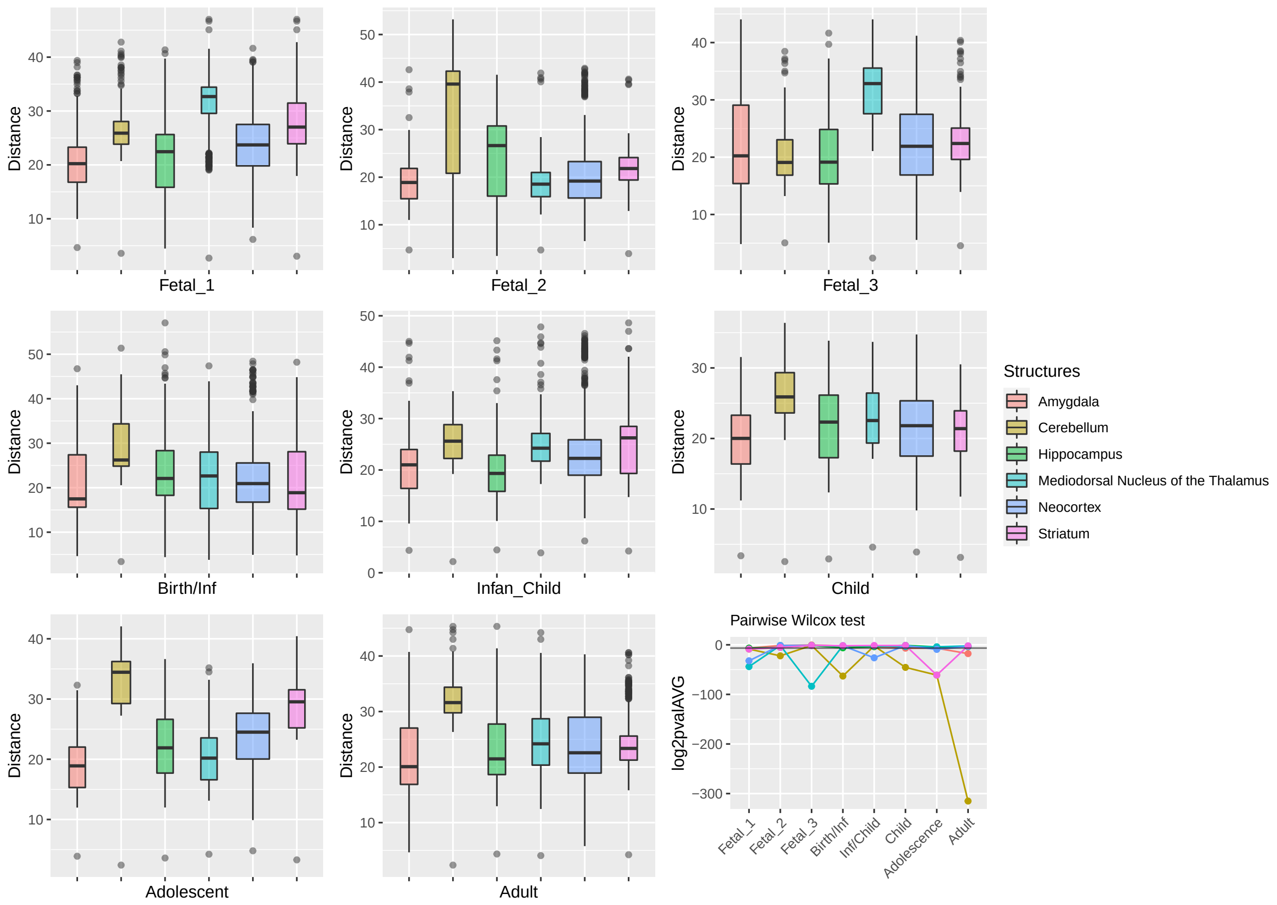


Figure 6: **Expression profile: Genes under positive selection not within large deserts of introgression.** Similarly to what is found in Supplementary Figure3 and 4, the cerebellum stand out postnatally (birth $p = 1.07\text{x}10^{-19}$, childhood $p = 1.95\text{x}10^{-14}$, adolescence $p = 4.11\text{x}10^{-19}$ and adulthood $p = 1.42\text{x}10^{-95}$). Other significant results are found for the thalamus (fetal stages 1 and 3). striatum (adolescence) or neocortex (fetal stage 1).

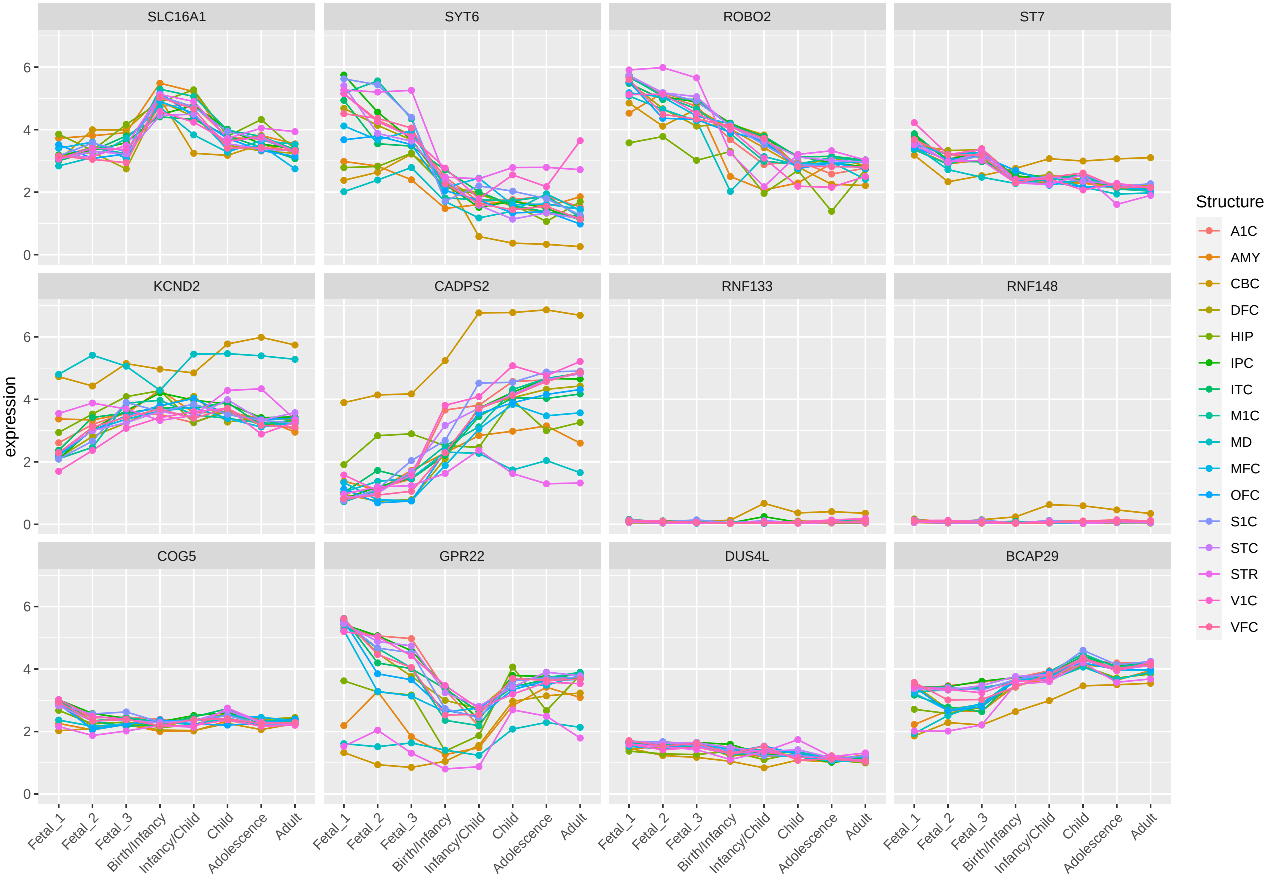


Figure 7: **Expression profile of genes within large deserts and positively-selected overlapping regions.** Twelve protein-coding genes were found at the intersection between large deserts of introgression and putative positively-selected regions. The expression of *CADPS2* and *KCND2* largely recapitulate the unique profile shown before (Figure S1) for the cerebellar cortex: Increasing expression from prenatal to postnatal stages, reaching the highest median expression value from childhood to adulthood in comparison to all other structures.
A1C, primary auditory cortex; AMY, amygdala; CBC, cerebellar cortex; DFC, dorsolateral prefrontal cortex; HIP, hippocampus; IPC, posterior inferior parietal cortex; ITC, inferior temporal cortex; M1C, primary motor cortex; MD, mediodorsal nucleus of thalamus; MFC, medial prefrontal cortex; OFC, orbital prefrontal cortex; S1C, primary somatosensory cortex; STC, superior temporal cortex; STR, striatum; V1C, primary visual cortex; VFC, ventrolateral prefrontal cortex.

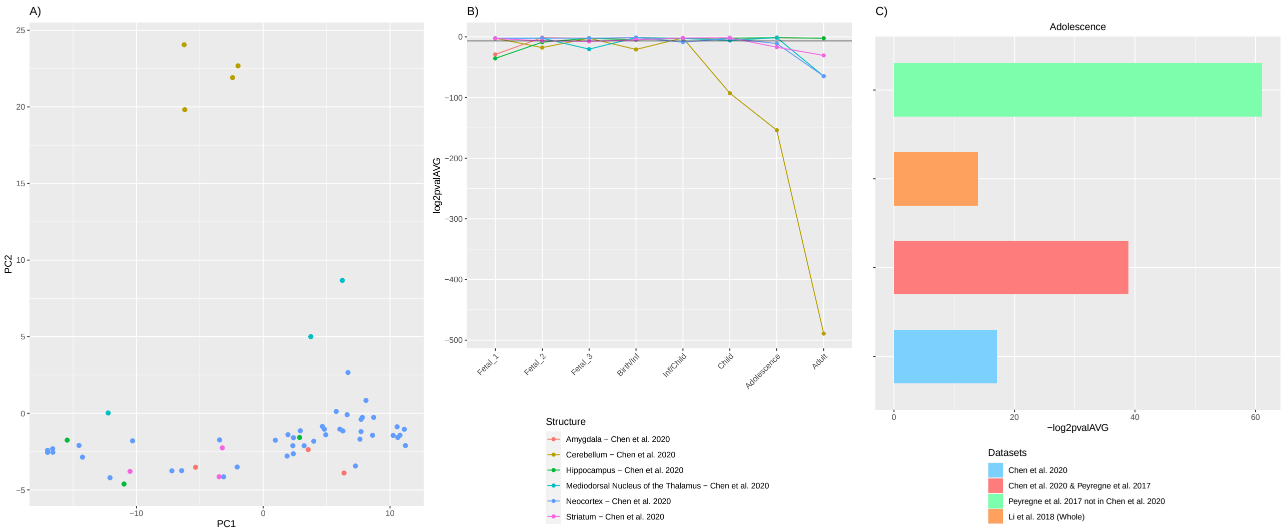


Figure 8: **The striatum's transcriptomic profile significantly diverges at adolescence for genes within large deserts.** When evaluating the transcriptome of genes within large deserts of introgression, the striatum reports significant differences postnatally at adolescence and adult stages. A) Distribution of structures at the adolescence using the first two principal components. B) P-values (log2-transformed) obtained from pairwise comparisons among structures at each developmental stage (Wilcoxon rank sum test with Bonferroni correction). C) Contribution of genes within large deserts of introgression [8], deserts of introgression under putative positive selection [8, 27], regions under putative selection [27] not within large deserts, and the raw dataset used in this study [1], to the observed divergence at adolescence for the striatum. The greatest value is found for genes under putative positive selection not within deserts of introgression, an effect also shown in Supplementary Figure 6.

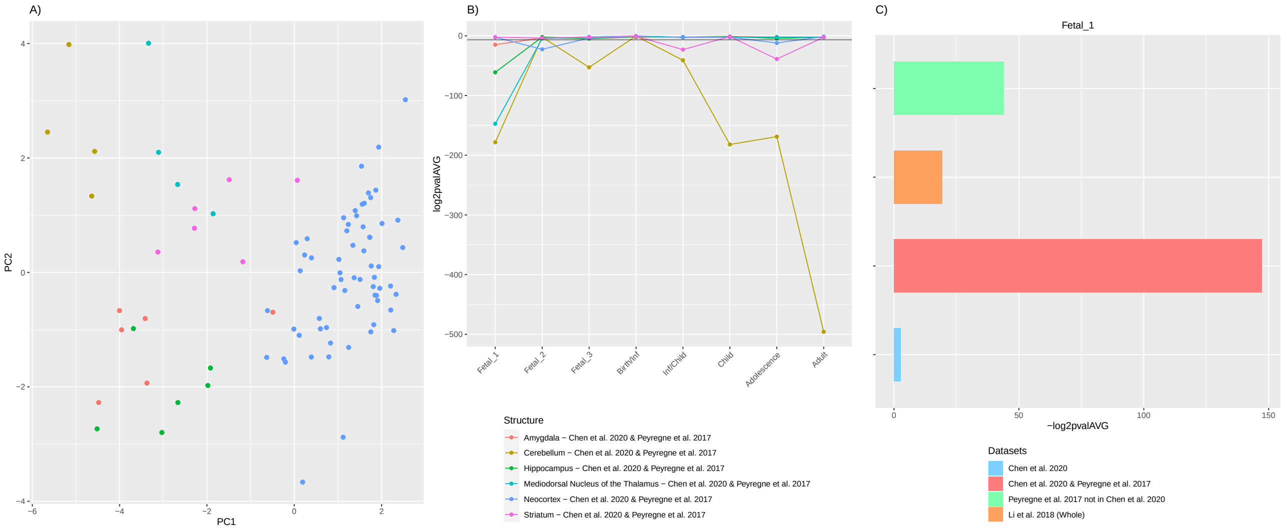


Figure 9: **The transcriptomic profile of the mediodorsal nucleus of the thalamus significantly diverges at fetal stage 1 for genes within deserts under positive selection.** A) Distribution of structures at the fetal stage 1 using the first two principal components. B) P-values (log2-transformed) obtained from pairwise comparisons among structures at each developmental stage (Wilcoxon rank sum test with Bonferroni correction). C) Contribution of genes within deserts of introgression [8], deserts of introgression under putative positive selection [8, 27], regions under putative selection [27] not within large deserts, and the raw dataset used in this study [1], to the observed divergence at adolescence for the striatum. The greatest value is found for genes within deserts under putative positive selection.

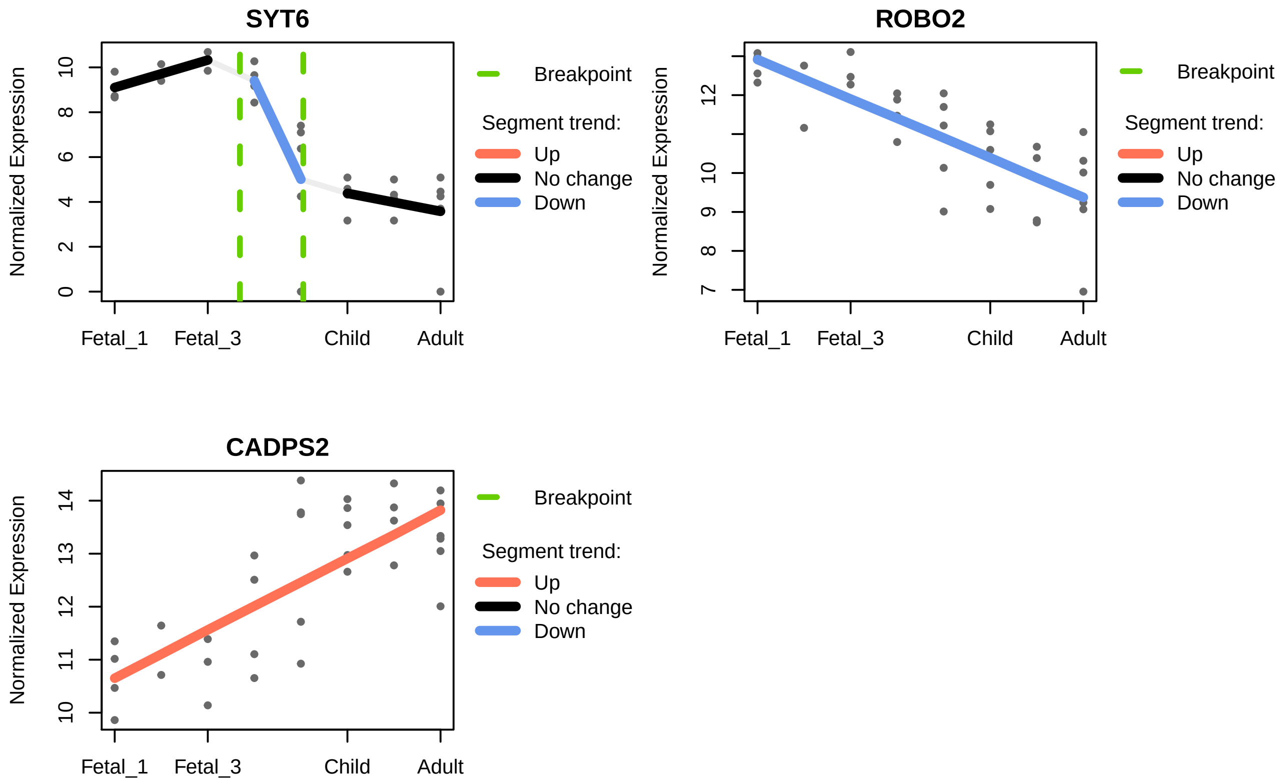


Figure 10: **Gene-specific trajectories - Cerebellar cortex.** A segmented regression analysis was performed to follow the trajectory of genes across developmental stages within both desertic and positively-selected regions. Only genes with an adjusted $R^2$ above 0.5 are shown.

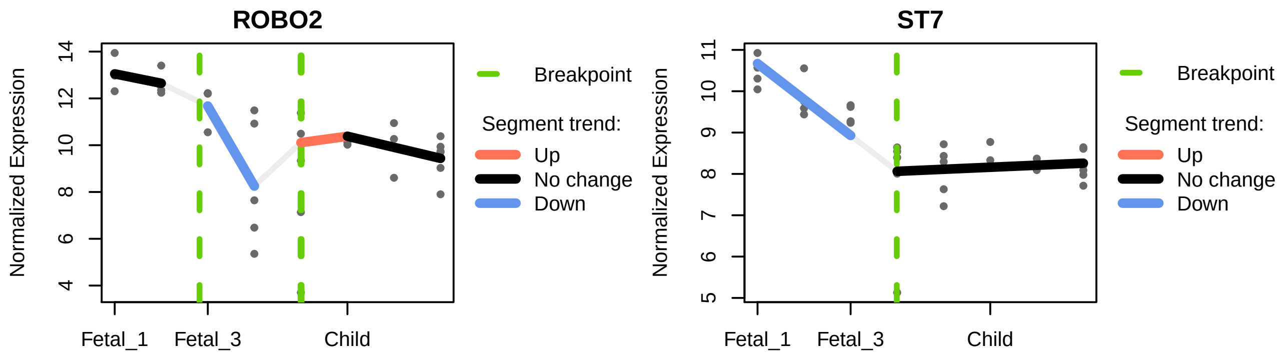


Figure 11: **Gene-specific trajectories - Thalamus.** Only genes within both desertic and positively-selected regions and an adjusted $R^2$ above 0.5 are shown.

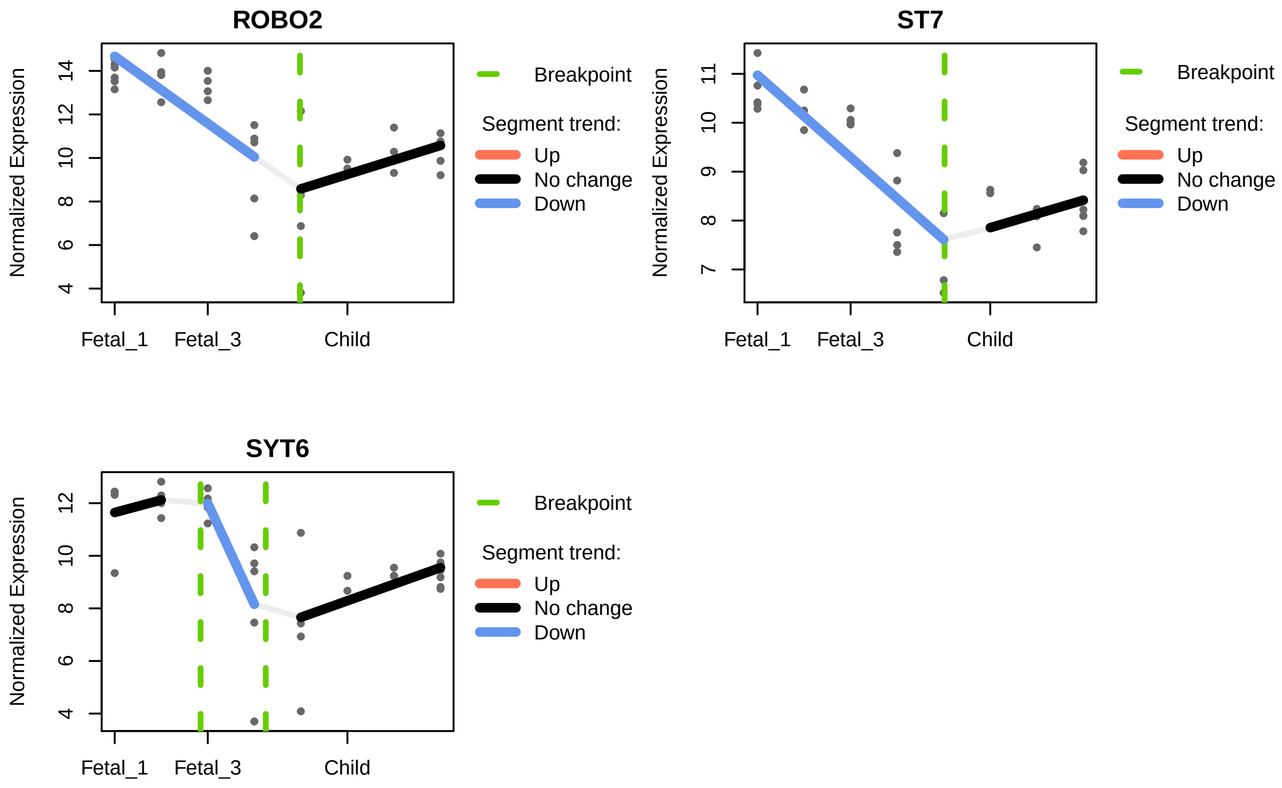


Figure 12: **Gene-specific trajectories - Striatum.** Only genes within both desertic and positively-selected regions and an adjusted $R^2$ above 0.5 are shown.
